# BodyMap transcriptomes reveal unique circular RNA features across tissue types and developmental stages

**DOI:** 10.1101/370718

**Authors:** Tong Zhou, Xueying Xie, Musheng Li, Junchao Shi, Jin J. Zhou, Kenneth S. Knox, Ting Wang, Qi Chen, Wanjun Gu

## Abstract

Circular RNAs (circRNAs) are a novel class of regulatory RNAs. Here, we present a comprehensive investigation of circRNA expression profiles across 11 tissues and 4 developmental stages in rats, along with cross-species analyses in humans and mice. Although positively correlated, circRNAs exhibit higher tissue specificity than cognate mRNAs. Also, genes with higher expression levels exhibit a larger fraction of spliced circular transcripts than their linear counterparts. Intriguingly, while we observed a monotonic increase of circRNA abundance with age in the rat brain, we further discovered a dynamic, age-dependent pattern of circRNA expression in the testes that is characterized by a dramatic increase with advancing stages of sexual maturity and a decrease with aging. The age-sensitive testicular circRNAs are highly associated with spermatogenesis, independent of cognate mRNA expression. The tissue/age implications of circRNAs suggest that they present unique physiological functions rather than simply occurring as occasional by-products of gene transcription.

## Introduction

Circular RNAs (circRNAs) are a class of endogenous RNAs with closed loop structures (Jeck and Sharpless, 2014). Many studies have revealed abundant circRNAs in organisms across the eukaryotic tree of life (Jeck et al., 2013; Wang et al., 2014). CircRNAs are mainly formed through pre-mRNA back-splicing (Jeck and Sharpless, 2014), which can be regulated by factors such as flanking intronic sequences (Zhang et al., 2014), RNA-binding proteins (Conn et al., 2015), canonical RNA splicing signals (Starke et al., 2015), and exon-containing lariat precursors (Barrett et al., 2015). CircRNAs have been present various biological functions, including acting as miRNA sponges (Memczak et al., 2013), transcriptional regulation of their parental genes (Li et al., 2015b), and RNA splicing regulation of their cognate messenger RNA (mRNA) (Conn et al., 2017). Moreover, several recent studies have provided strong evidence that some circRNAs can be translated in a cap-independent manner (Legnini et al., 2017; Pamudurti et al., 2017; Yang et al., 2017). The expanding view of circRNA biogenesis and function suggests that circRNAs are a novel class of RNAs with important biological implications.

Given the regulatory functions of circRNAs in gene expression, investigation of the dynamic expression of circRNAs across different cell types, tissues, and organisms is helpful for understanding their roles in various biological processes. CircRNAs are conservatively expressed from orthologous genomic regions between humans and mice (Guo et al., 2014; Jeck et al., 2013) and among *Drosophila* species (Westholm et al., 2014). Detailed analyses of circRNA expression in humans (Guo et al., 2014; Liu et al., 2016b; Xia et al., 2016), mice (Xia et al., 2016), pigs (Liang et al., 2017b), and flies (Westholm et al., 2014) have revealed substantial tissue-specific patterns of circRNA expression. Notably, circRNAs are enriched and abundantly expressed in some specific tissue types and blood components, such as the brain (Rybak-Wolf et al., 2015; Szabo et al., 2015; Veno et al., 2015; Westholm et al., 2014), testes (Liang et al., 2017b), peripheral whole blood (Memczak et al., 2015), peripheral blood mononucleotide cells (Qian et al., 2017), platelets (Alhasan et al., 2016), and exosomes (Li et al., 2015a). CircRNAs have also been related to the development of the fetal human brain (Szabo et al., 2015), mouse brain (You et al., 2015), fetal porcine brain (Veno et al., 2015), and *Drosophila* neural systems (Westholm et al., 2014). Interestingly, neural expression of some circRNAs in flies has been suggested as a potential biomarker of aging (Westholm et al., 2014). In addition, aberrant circRNA expression is related to human diseases (Chen et al., 2016), including human cancers (Meng et al., 2017), neural degenerative diseases (Kumar et al., 2016), hematological malignancies (Bonizzato et al., 2016), and infectious diseases (Qian et al., 2017).

Although substantial advances have been made in understanding circRNA expression and its potential function, little is known about the correlation between the expression of circRNAs and mRNAs transcribed from the same host genes. Since circRNAs can either promote the transcription of their host gene (Li et al., 2015b) or regulate the splicing of cognate mRNAs (Conn et al., 2017), the correlation between circRNAs and their linear counterparts can be dynamically regulated. Therefore, it is important to obtain an in-depth understanding of the relationship between the expression profiles of circRNAs and their cognate mRNAs across tissue types and developmental stages. Some previous studies (Chen, 2016; Conn et al., 2015; Guo et al., 2014; Liang and Wilusz, 2014; Rybak-Wolf et al., 2015; You et al., 2015) have shown that there is no clear correlation between the expression values of circRNAs and their corresponding mRNAs. However, these conclusions might be preliminary and premature, since the circRNA profiles examined in these studies were derived from relatively small datasets, such as RNA sequencing (RNA-seq) data from a single tissue type (Conn et al., 2015; Rybak-Wolf et al., 2015; You et al., 2015), the results of mini-gene experiments (Liang and Wilusz, 2014), or data from several different tissues and cell types from different publications (Guo et al., 2014). Since a batch effect inevitably exists in high-throughput sequencing data collected from various sources (Leek et al., 2010), it is technically difficult to conduct a comprehensive comparison of mRNA and circRNA expression across tissues or developmental stages. In addition, the computational estimation of expression values for both linear and circular transcripts based on rRNA-depleted RNA-seq data may not be optimized (Gao and Zhao, 2018). The first reason for this lack of optimization is that circRNA expression values in previous studies (Conn et al., 2015; Guo et al., 2014; Rybak-Wolf et al., 2015; You et al., 2015) have been quantified based on the ratio of back-splicing reads to canonical linear reads at a given junction from RNA-seq data. However, this count-based quantification method is less accurate than model-based approaches (Kanitz et al., 2015). The second reason is that canonical reads corresponding to circular transcripts could be mis-assigned with their corresponding linear transcripts using classical RNA-seq quantification tools. Therefore, it is important to consider both circular and linear transcripts when quantifying RNA expression values from RNA-seq data.

To overcome above issues, we analyzed the transcriptomes of both mRNAs and circRNAs in a rat BodyMap RNA-seq dataset (Yu et al., 2014a; Yu et al., 2014b) using *Sailfish-cir* (Li et al., 2017), a computational tool that we recently developed, which applies a model-based algorithm to precisely quantify expression level of both linear and circular transcripts from rRNA-depleted RNA-seq data. The rat BodyMap dataset contains 320 samples isolated from *Fischer 344* rats across 11 tissues and 4 developmental ages (Yu et al., 2014a; Yu et al., 2014b). This is an ideal dataset for a systematic comparison of expression values of circRNAs and its cognate mRNAs across tissue types and developmental stages. Using the rat BodyMap dataset, we compiled a repertoire of circRNAs in the rat transcriptome, and summarized the expression profiles of both circRNAs and mRNAs in all these 320 samples. We explored the expression patterns of circRNAs and linear mRNAs across 11 rat tissues and 4 developmental stages, and investigated the relationship in expression between circRNAs and their linear counterparts. Furthermore, we looked into the roles of circRNAs in determining the tissue’s phenotypes and their relations to rat development, and we compared their biological implications with those of linear mRNAs. We found that i) circRNAs are evolutionarily more conserved than mRNAs; ii) although the expression of circRNAs is positively correlated with that of cognate mRNAs, genes with higher expression levels tend to show a significantly larger fraction of spliced circular transcripts than their linear counterparts; iii) circRNAs exhibit higher tissue specificity than mRNAs; iv) circRNA abundance monotonically increases with age in the rat brain, which is consistent with observations made in humans (Szabo et al., 2015), mice (Gruner et al., 2016; You et al., 2015), and flies (Westholm et al., 2014); and v) testes circRNA expression shows a dynamic age-dependent pattern, with a dramatic increase with advancing stages of sexual maturity (2, 6, and 21 weeks), followed by a decrease with aging (104 weeks). The age-sensitive testicular circRNAs are highly associated with spermatogenesis (e.g., cilium morphogenesis and spermatid development) and independent of the expression of their cognate mRNAs. Our study elucidates a complex landscape of circRNA expression across different tissues and developmental stages, and thorough comparison of circRNA expression profiles against their linear counterparts provides us with a deeper understanding of the biological roles of circular transcripts in tissue specificity and development as well as their relationship with linear mRNAs.

## Results

### A comprehensive rat circRNA repertoire

To analyze the expression profiles of circRNAs in the rat BodyMap dataset, a reference library of known rat circRNAs is required (see *Methods* for details), but no such library is available in public databases (Glazar et al., 2014; Liu et al., 2016a). Therefore, we first constructed a repertoire of rat circRNAs from 320 samples. Because it is difficult to distinguish circRNA isoforms that are generated by the same back-splicing event using short-read RNA-seq data (Gao et al., 2016), all circRNA isoforms with the same back-splicing junction were considered “circRNA species.” The subsequent analyses described in this section were performed at the circRNA species level.

Using our computational pipeline, a total of 16,745 circRNA species were identified in the rat BodyMap dataset. Among different rat tissues, the brain expressed the greatest number of circRNA species, while the liver expressed the fewest circRNA species (Figure 1A). The majority (86.6%, 14,493 out of 16,745) of rat circRNA species were derived from exonic regions (Figure 1B). In comparison, only 1,223 (7.3%) and 1,029 (6.1%) circRNA species originated from intronic and intergenic regions, respectively (Figure 1B). Among the exonic circRNAs, most were composed of less than 5 exons, although some contained more than 20 exons (Figure 1C). As expected, the length of exonic circRNAs was significantly and positively correlated with the number of exons (*Spearman*’s rank correlation test: *P* < 10^-10^; Supplementary Figure S1), and most rat exonic circRNAs were less than 1,000 base pairs in length (Figure 1D). Given that exonic circRNAs were predominant in the rat circRNA repertoire, we focused on only exonic circRNAs in the rest of this study.

**Figure 1.**
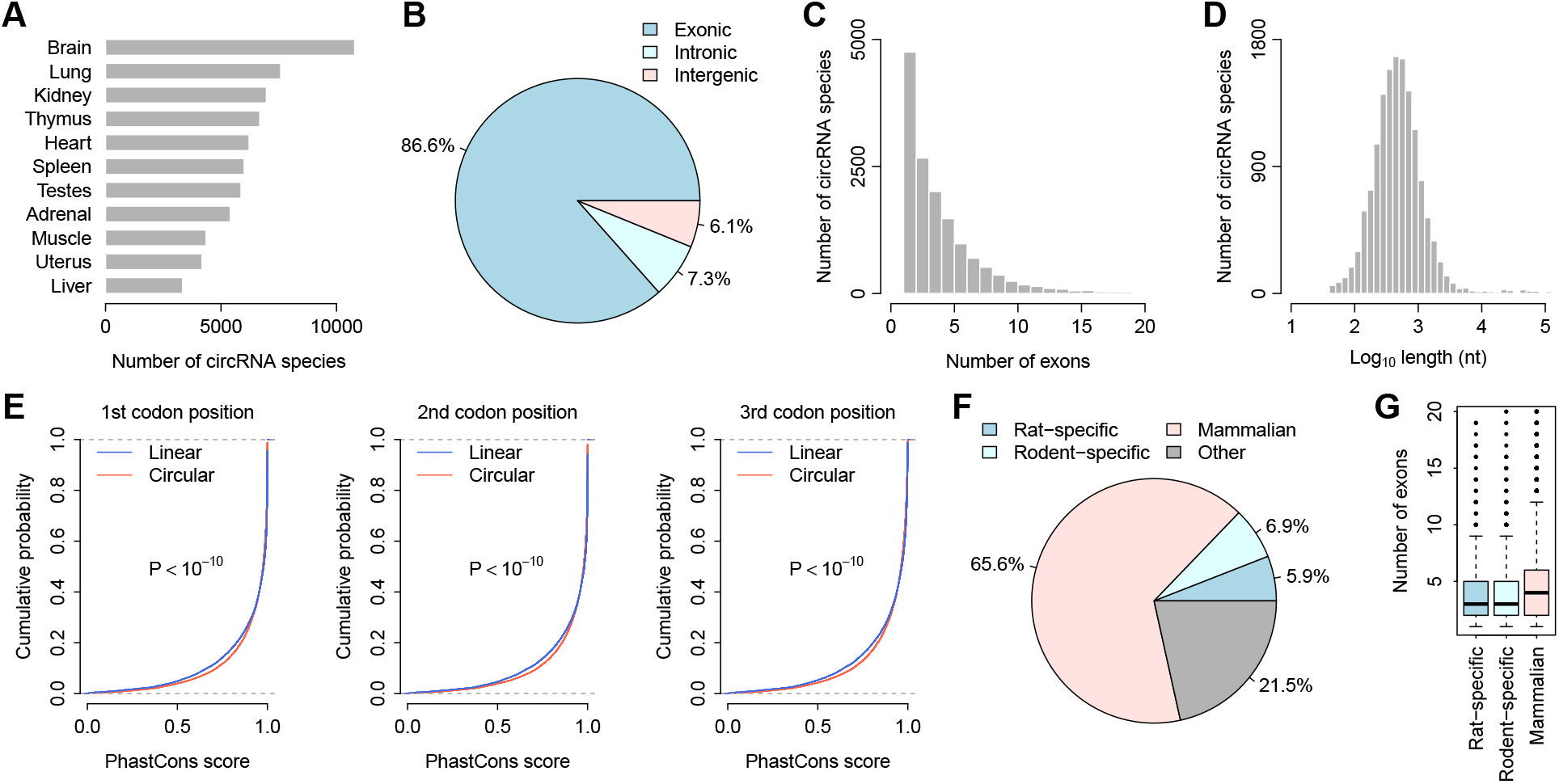
The rat circRNA repertoire. (A) Comparison of the number of circRNA species across the 11 rat tissues. (B) Fractions of exonic, intronic, and intergenic circRNA species within the repertoire. (C) Histogram of the number of exons in the exonic circRNAs of rats. (D) Histogram of circRNA length in rats. (E) Cumulative distribution of the *PhastCons* scores of both linear and circular transcripts at the three codon positions. A significantly increased *PhastCons* score was observed for the circular transcripts compared with the linear transcripts. The P-values were calculated with the *Kolmogorov-Smirnov* test. (F) Fractions of mammalian-common, rodent-specific, and rat-specific circRNA species within the repertoire. (G) Comparisons of exon numbers among the mammalian-common, rodent-specific, and rat-specific circRNAs.

We next analyzed the evolutionary conservation level (i.e., *PhastCons* score) (Pollard et al., 2010) at the three codon positions in both circRNA and mRNA exons. We found that the conservation level of exonic circRNAs was significantly higher than that of mRNAs (*Kolmogorov-Smirnov* test: *P* < 10^-10^ for all three codon positions; Figure 1E). This finding suggests that circRNAs may be under some extra selective pressure to ensure their proper biogenesis and/or functions, such as binding of splicing factors or RNA-binding proteins. Furthermore, we investigated the evolutionary dynamics of circRNA biogenesis in the mammalian lineage. We compared the rat circRNA repertoire with the human and mouse circRNA repertoires. We observed that most of the rat exonic circRNA species occurred in all three mammalian transcriptomes (Figure 1F). In comparison, only 997 (6.9%) of the rat exonic circRNAs evolved recently, after the separation of the common ancestor of rodents and humans (Figure 1F). Additionally, 861 (5.9%) of the rat exonic circRNAs evolved specifically in the rat lineage (Figure 1F). These rat-specific and rodent-specific circRNAs contained fewer exons than the mammalian-common circRNAs (*Wilcoxon* test: *P* < 10^-10^; Figure 1G), which suggests that younger circRNAs are more likely to be short in length. Given the importance of flanking RNA structure in circRNA biogenesis (Ashwal-Fluss et al., 2014; Liang and Wilusz, 2014; Zhang et al., 2014), it is reasonable to observe relatively shorter circular transcripts in evolutionarily younger circRNA species.

### Relationship between the expression of circular and linear transcripts

Based on the rat circRNA repertoire described above, we estimated the expression levels of both circular and linear transcripts in all rat BodyMap samples using our recently published RNA-seq quantification framework, *Sailfish-cir* (Li et al., 2017). For each host gene, the transcripts per million (*TPM*) values of both circular (*TPM_circ_*) and linear transcripts (*TPM_linear_*) were calculated. We observed a significant positive correlation between the expression of circRNAs and their linear counterparts in all 11 tissue types (*Spearman*’s rank correlation test: *P* < 10^-10^; Figures 2A and 2B and Supplementary Figure S2), which suggests that circRNA expression is largely regulated at the transcriptional level of the corresponding host genes. Across the 11 tissue types, the expression of circRNAs was significantly lower than that of their linear counterparts (paired *t*-test: *P* < 10^-10^). However, there were a considerable number of circRNAs with relatively high expression levels, exceeding those of their linear counterparts (Figures 2A and Supplementary Figure S2). For example, *TPM_circ_* was found to be higher than *TPM_linear_* for 462 host genes in the rat brain (Figure 2A). We also calculated the fraction of circular transcripts [*TPM_circ_* / (*TPM_circ_* + *TPM_linear_*)] for individual host genes. Interestingly, we found that, across the 11 tissue types, genes with higher expression levels tended to exhibit a significantly larger fraction of spliced circular transcripts than their linear counterparts (*Spearman*’s rank correlation test: *P* < 10^-10^; Figure 2C), which suggests that circRNAs are more “sensitive” to alterations of host gene expression than mRNAs. This observation also implies that in addition to being subject to transcriptional regulation of host genes, the expression level of circRNAs may also be controlled at the splicing level.

**Figure 2.**
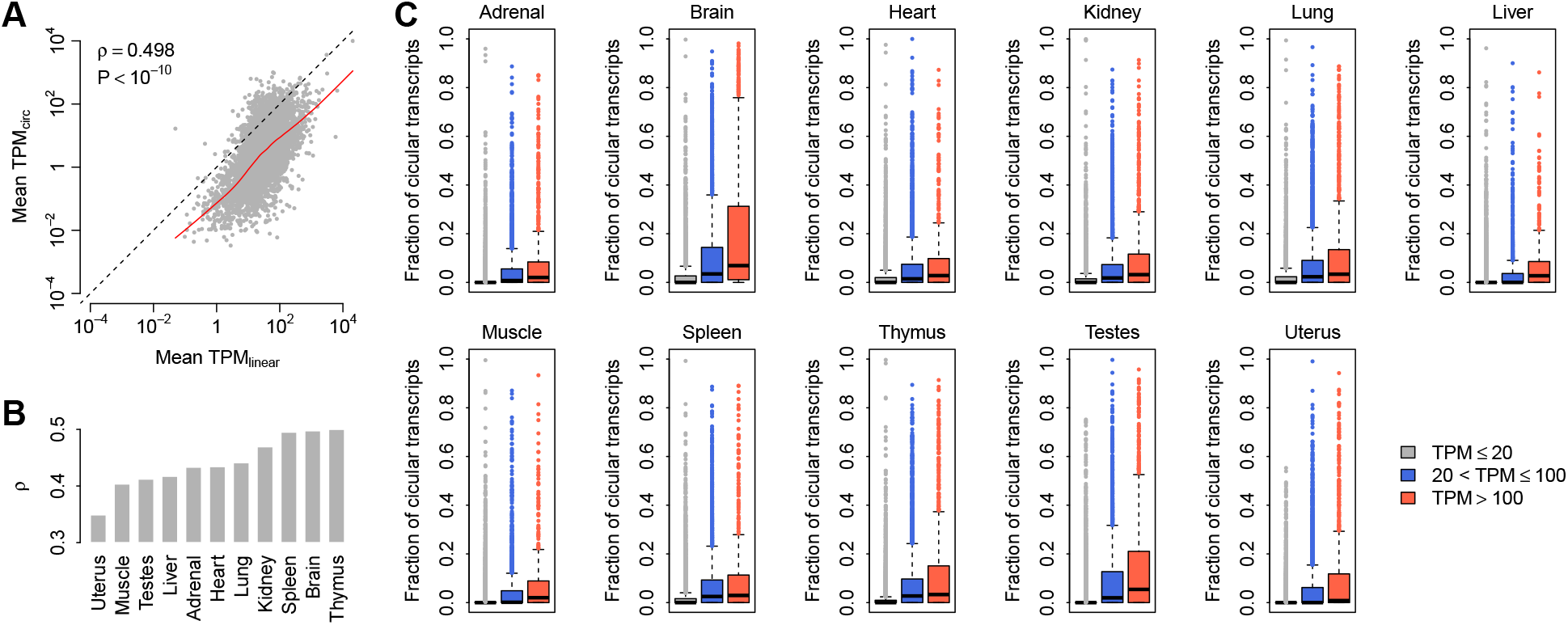
Relationship between the expression of circRNAs and their linear counterparts. (A) Correlation between mean *TPM_linear_* and mean *TPM_circ_* values in the rat brain. Each dot represents one host gene. The dots above the diagonal denote the host genes with higher circRNA expression relative to their linear transcripts in the brain. The red curve represents *Lowess* smoothed data. The correlation coefficient (*ρ*) and *P*-value were calculated with Spearmans rank correlation test. (B) Comparison of the correlation coecient (*ρ*) calculated between mean *TPM_linear_* and mean *TPM_circ_* values across the 11 rat tissues by *Spearman’s* rank correlation test. (C) Comparison of the fraction of circular transcripts for genes categorized by expression level.

### CircRNAs show higher tissue specificity than mRNAs

Tissue-specific expression has been systematically investigated for mRNAs but not circRNAs (Andergassen et al., 2017; Fagerberg et al., 2014; Yu et al., 2014a). To understand the extent to which circRNA expression shows a tissue-dependent pattern, we performed a principal component analysis (PCA) of circRNA expression for all the rat tissue samples. We found that the samples from the same tissue type tended to cluster together according to the first and second principal components (Figure 3A), which suggests that circRNAs are expressed in a tissue-specific manner. Notably, the brain and testes samples showed extremely unique PCA patterns compared with the other tissues (Figure 3A), which was further confirmed by the observation that the brain and testes exhibited the largest number of highly expressed circRNAs (Figure 3B). Additionally, the mean *TPM_circ_* values and the fraction of circular transcripts were significantly higher in the brain and testes than in the other tissue types (*Kolmogorov-Smirnov* test: *P* < 10^-5^; Figures 3C and 3D).

**Figure 3.**
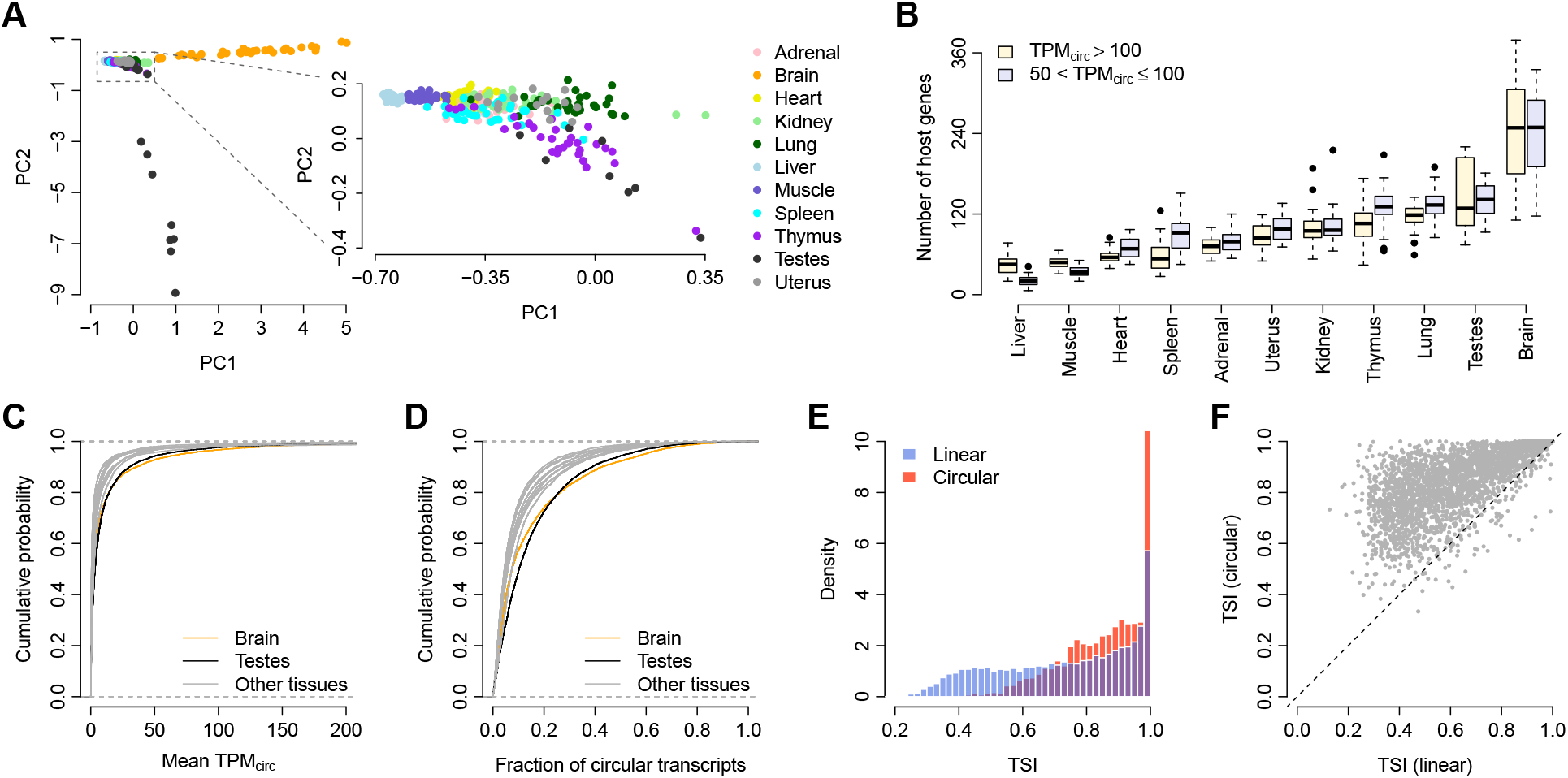
Tissue specicity of circRNA expression. (A) PCA of circRNA expression. Each dot represents one tissue sample. PC1: the first principal component; PC2: the second principal component. (B) Number of host genes with high circRNA expression. Genes with relatively higher *TPM_circ_* were categorized into two groups: *TPM_circ_* > 100 and 50 < *TPM_circ_* < 100. Each boxplot consists of 32 samples from the specific tissue type, except for the testes and uterus, for which only 16 samples are included. (C) Cumulative distribution of the mean *TPM_circ_* values across all 11 tissue types. (D) Cumulative distribution of the fraction of circular transcripts across all 11 tissue types. (E) Histogram of the *TSI* of both linear and circular transcripts. (F) Paired comparison of *TSI* between linear and circular transcripts. Each dot represents one host gene. The dots above the diagonal denote the host genes with a higher circRNA *TSI* than their linear counterparts.

To compare tissue specificity between circRNAs and mRNAs, we calculated the tissue specificity index (*TSI*) for both the circular and linear transcripts of each host gene (see *Methods* for details). A higher *TSI* indicates higher tissue specificity. We found that there were more circRNAs than linear RNAs showing a *TSI* > 0.8 (Figure 3E). Paired comparisons indicated that the *TSI* of circular transcripts was significantly higher than the *TSI* of their corresponding linear counterparts (paired *Wilcoxon* test: *P* < 10^-10^; Figure 3F). All these results suggest that, although mRNAs exhibit a tissue-specific expression pattern (Supplementary Figure S3), circRNAs are expressed in a more dynamic manner among different rat tissues and show higher tissue specificity.

### A map of tissue-specific circRNAs and their potential physiological functions

To understand whether the observed tissue-specific circRNA expression is relevant to the physiological function of the specific tissue, we performed a hierarchical clustering analysis based on the dynamic expression of the tissue-specific circRNAs across 320 rat tissue samples. We observed that the samples from the same tissue type were clustered into a single group, and each tissue type exhibited one or more unique tissue-specific circRNA block(s) (Figure 4A). Tissues with similar physiological functions tended to group together and share common circRNA blocks, as observed for the thymus and spleen, which play vital roles in the immune system (Figure 4A). Additionally, the samples from two reproduction-related tissues, testes and uterus, were aggregated into one group as well (Figure 4A). In contrast, the brain samples showed patterns distinct from those of other tissues (Figure 4A).

**Figure 4.**
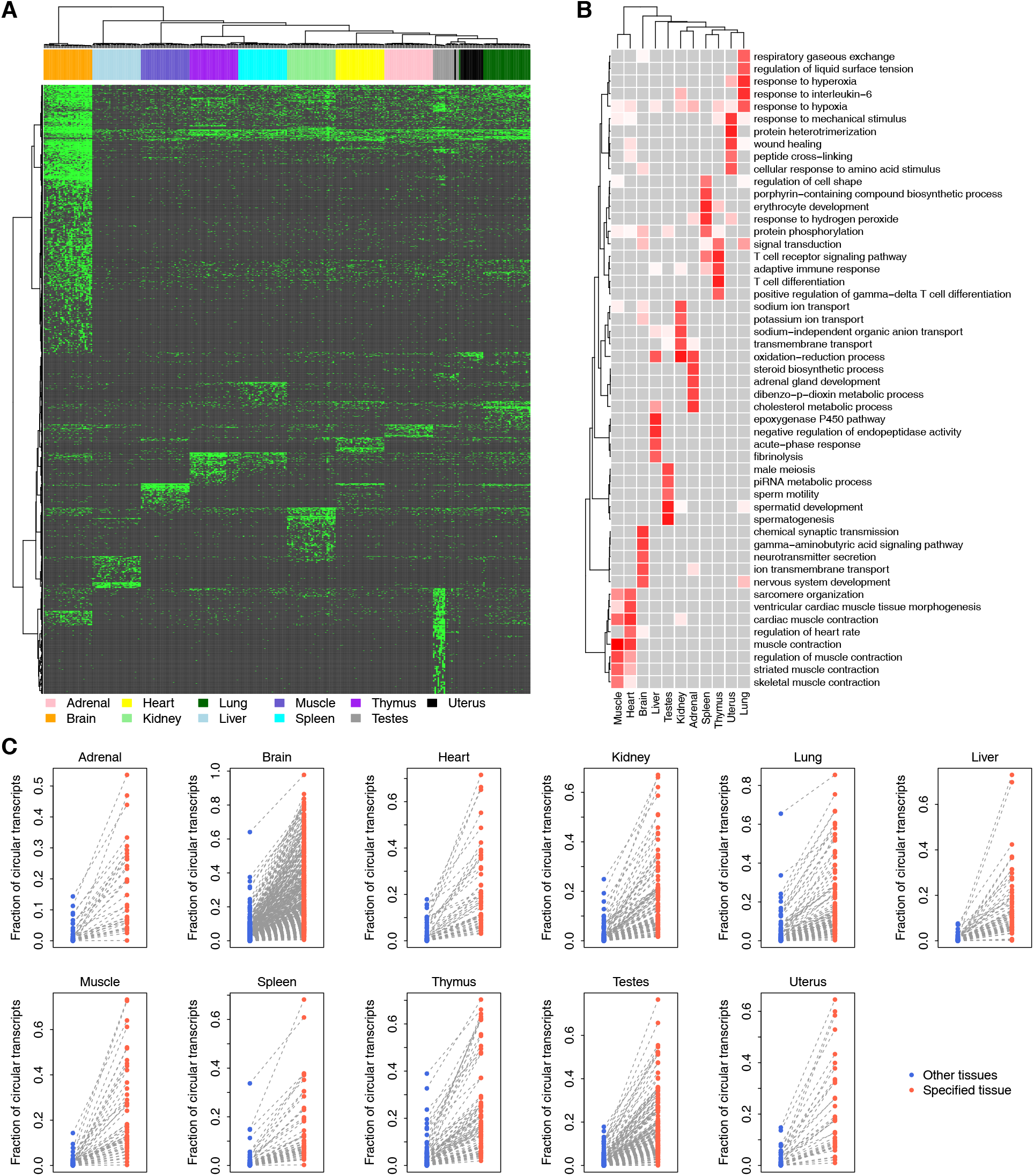
Map of tissue-specific circRNAs. (A) Hierarchical clustering of tissue-specific circRNAs. Each green dot represents one expressed tissue-specific circRNA. (B) The top GOBP terms associated with the host genes of the tissue-specific circRNAs. For each tissue type, the top five GOBP terms are listed. The association between the GOBP terms and tissue-specific circRNAs was measured based on the *Z*-score calculated from *Fisher’s* exact test. Darker red indicates a stronger association, while lighter red indicates a weaker association. Grey indicates no association. (C) Fraction of circular transcripts of tissue-specific circRNAs. Each dot represents one tissue-specific circRNA.

To explore the potential functions of circRNAs in different tissues, we next examined the Gene Ontology Biological Process (Ashburner et al., 2000) (GOBP) terms associated with the host genes of the tissue-specific circRNAs. We found that the enriched GOBP terms were highly related to the biological/physiological function of each specific tissue (Figure 4B). For example, the lung-specific circRNAs were associated with “respiratory gaseous exchange”, “response to hyperoxia”, and “response to hypoxia”; the brain-specific circRNAs were associated with “chemical synaptic transmission”, “neurotransmitter secretion”, and “nervous system development”; and the testes-specific circRNAs were associated with “sperm motility”, “spermatid development”, and “spermatogenesis” (Figure 4B). All these results suggest that the tissue-specific expression pattern of circRNAs is important for all types of tissues to perform their biological/physiological roles.

To gain further insight into the potential driving force causing the tissue-specific expression of circRNAs, we compared the splicing ratio of tissue-specific circRNAs in the specific tissue against that in all other tissue types. We observed a consistently higher fraction of circular transcripts for the tissue-specific circRNAs in an individual tissue (paired *Wilcoxon* test: *P* < 10^-10^; Figure 4C). However, we observed an opposite pattern when tissue-specific mRNAs were considered (Supplementary Figure S4). These results suggest that tissue-specific circRNAs, but not tissue-specific linear mRNAs, are positively regulated at the splicing or post-transcriptional level to ensure the proper function of the specific tissue.

### Age-dependent circRNA expression in the rat brain

We next investigated the temporal changes in circRNA expression in rat tissues at four different developmental stages (2, 6, 21, and 104 weeks). Although the overall fraction of circular transcripts was not longitudinally correlated with age for most of the tissues (Figure 5A), the abundance of brain circRNAs was found to monotonically increase with age (*Spearman*’s rank correlation test: *ρ* = 0.772 and *P* = 2.3×10^-7^; Figure 5A), which is consistent with previous observations made in the brains of humans (Szabo et al., 2015), mice (Gruner et al., 2016; You et al., 2015), and flies (Westholm et al., 2014). We also investigated the age-dependent expression of each gene in the brain. The correlation coefficient (*ρ*) between expression and age was calculated for both circular and linear transcripts. We found that the *ρ* values of the circRNAs were significantly more positive than those of the linear RNAs (*t*-test: *P* < 10^-10^; Figure 5B). In particular, when we focused on the circRNAs with a strong positive correlation (*ρ* > 0.7), we found that the *ρ* values of their corresponding linear counterparts were significantly lower (paired *t*-test: *P* < 10^-10^; Figure 5C). All these results suggest that the age-dependent expression of brain circRNAs is largely independent of linear transcripts and, at least to a certain extent, is not a secondary consequence of host gene expression. Gene ontology analysis indicated that the circRNAs that accumulated with age were enriched in GOBP terms linked to neural development, such as “brain-derived neurotrophic factor receptor signaling pathway” and “activation of GTPase activity” (Figure 5D). Our data support the concept that circRNAs in the brain might play an essential role in regulating synaptic plasticity and neuronal differentiation (Hanan et al., 2017).

**Figure 5.**
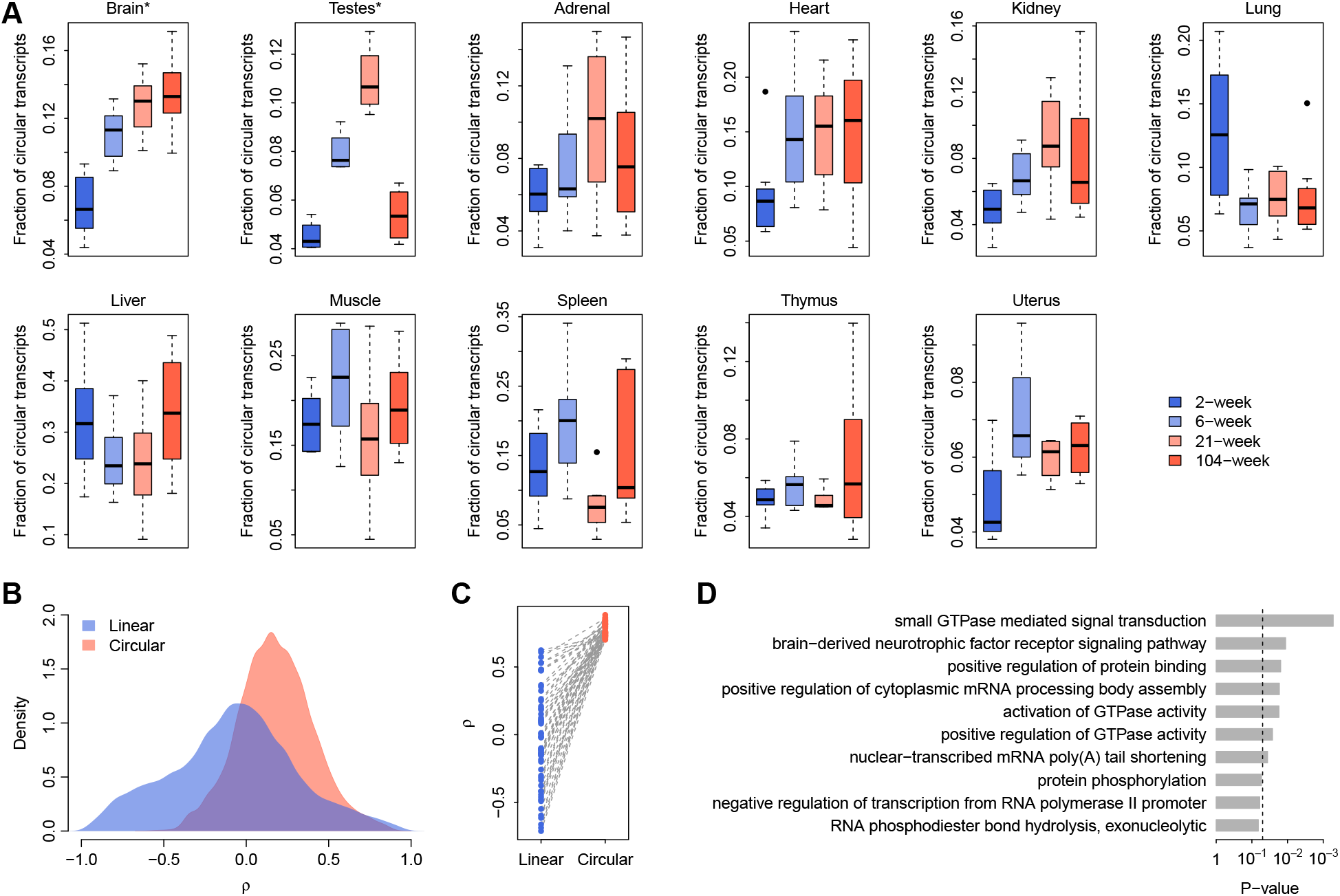
Age-dependent circRNA expression. (A) Relationship between age and overall circRNA abundance across the 11 tissue types. (B) Distribution of the correlation coefficients (*ρ*) between age and the expression of linear/circular transcripts in the brain. The *rho* values were calculated with *Spearman’s* rank correlation test. (C) Paired comparisons of *ρ* between the circRNAs that accumulated with age (*ρ* > 0:7) and their linear counterparts in the brain. (D) Top 10 GOBP terms associated with the circRNAs that accumulated with age in the brain. The *P*-values were calculated with *Fisher’s* exact test. The vertical dashed line indicates the significance level of *α* = 0.05.

### Age-dependent circRNA expression in testes

Interestingly, circRNA expression in rat testes showed a more dynamic pattern during the four developmental stages (2, 6, 21, and 104 weeks) we examined (Figure 5A). Unlike the monotonic increase of circRNA abundance in the rat brain, the overall fraction of circular transcripts in testes accumulated linearly for the first three developmental stages (2, 6, and 21 weeks), which nicely mirrored the stages approaching sexual maturity, during which the male reaches reproductive peak. However, the abundance of circRNAs drastically decreased at the age of 104 weeks (Figure 5A), at which stage the rats could be classified as aged males with declining reproduction. Moreover, by further examining the correlation between expression and developmental stage for individual genes, we showed that for the circRNAs exhibiting monotonically increased expression from 2 weeks to 21 weeks (*ρ* > 0.6), the *ρ* values of their corresponding linear counterparts were significantly lower (paired *t*-test: *P* < 10^-10^; Figure 6A). On the contrary, for the circRNAs downregulated from 21 weeks to 104 weeks (*ρ* <-0.6), the *ρ* values of their linear counterparts were significantly increased (paired *t*-test: *P* < 10^-10^; Figure 6A), which suggests that the age-dependent expression of testes circRNA is also somewhat independent of host gene expression and may represent a unique signature of the stage-specific reproductive performance of the male. Indeed, gene ontology analysis further revealed that both the genes upregulated from 2 weeks to 21 weeks and the genes downregulated from 21 weeks to 104 weeks were significantly enriched in GOBP terms linked to spermatogenesis, such as “cilium morphogenesis”, “spermatid development”, and “spermatogenesis” (Figure 6B).

**Figure 6.**
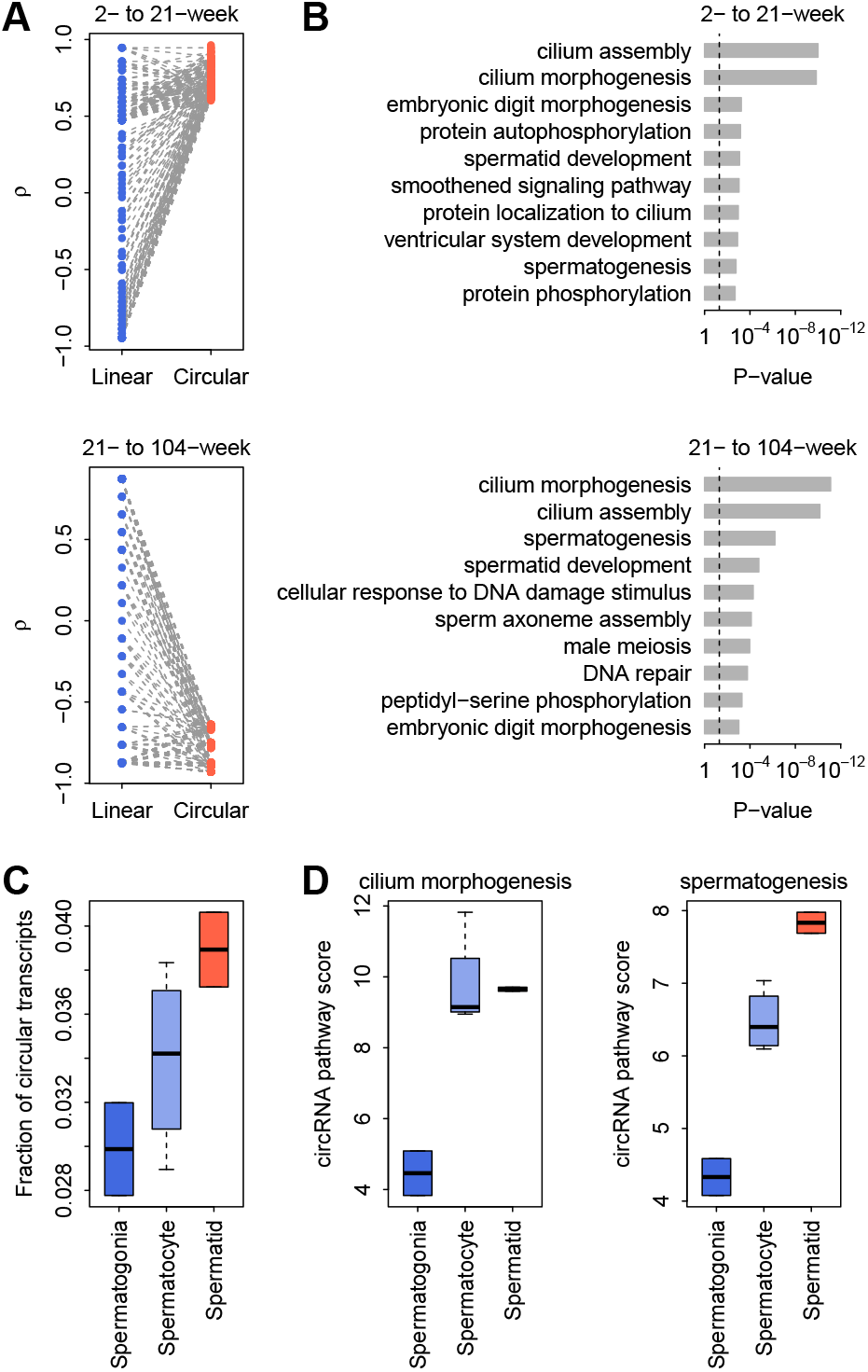
Age-dependent circRNAs in testes. (A) Paired comparisons of correlation coefficients (*ρ*) between the age-dependent circRNAs (*ρ* > 0:6) and their linear counterparts in rat testes. The *ρ* values were calculated between age and the expression of linear/circular transcripts using *Spearman’s* rank correlation test. (B) The top 10 GOBP terms associated with the age-dependent circRNAs in rat testes. The *P*-values were calculated with *Fisher’s* exact test. The vertical dashed line denotes the significance level of *α* = 0:05. (C) The circRNA abundance in mouse spermatogenic cells was categorized by distinct stages (spermatogonia, spermatocytes, and spermatids, sequentially). (D) The circRNA expression-based pathway scores of the mouse spermatogenic cells. We calculated the circRNA pathway scores for both the “cilium morphogenesis” and “spermatogenesis” GOBP terms. Higher pathway scores indicate higher overall circRNA expression for the specific GOBP terms.

The age-specific pattern of circRNAs found in rat testes might be correlated at the molecular level with certain stages of sperm development that are indicative of specific functions. To test this hypothesis, we further examined the stage-specific circRNA expression profiles of different spermatogenic cells (spermatogonia, spermatocyte, and spermatid) from a published dataset for mice (Lin et al., 2016) and cross-analyzed the spermatogenic stage-specific gene categories with the age-specific circRNA signatures observed in rats. Interestingly, we found that the overall fraction of circular transcripts increased monotonically with the spermatogenesis stage in mice (spermatogonia, spermatocytes, and spermatids, sequentially) (*Spearman*’s rank correlation test: *ρ* = 0.772 and *P* = 2.5×10^-2^; Figure 6C). We also found that the circRNA expression-based pathway scores (see *Methods* for details) for both the “cilium morphogenesis” and “spermatogenesis” GOBP terms (most dynamically associated with age in rat testes) were positively correlated with the spermatogenic stage in mice (*Spearman*’s rank correlation test: *ρ* = 0.772 and *P* = 2.5×10^-2^ for “cilium morphogenesis”, and *ρ* = 0.926 and *P* = 9.6×10^-4^ for “spermatogenesis”; Figure 6D). These results further support our hypothesis that the circRNAs present in testes not only are highly associated with the developmental stages of spermatogenesis but also could be harnessed as a biomarker for reproductive aging.

## Discussion

In this study, we performed a thorough investigation of circRNA transcriptomes in the rat BodyMap dataset. In comparison with existing studies (Conn et al., 2015; Guo et al., 2014; Rybak-Wolf et al., 2015; You et al., 2015), we systematically analyzed the expression profiles of both circular and linear transcripts in 320 rat samples across 11 tissue types and 4 developmental stages and particularly focused on the relationships and differences between the expression of circRNAs and mRNAs. Understanding the relationship between circRNAs and mRNAs is helpful for answering several open questions regarding circRNA biogenesis and function, including i) whether circRNAs are simply by-products of mRNA transcription, ii) whether circRNAs show expression patterns similar to those of mRNAs across different tissue types, and iii) whether the association of circRNAs with aging is similar to that of mRNAs.

First, we asked whether circRNAs are simply by-products of the transcription and splicing of their host genes. If not, what are the factors affecting the final products of circular transcripts? Several groups (Conn et al., 2015; Liang and Wilusz, 2014; Rybak-Wolf et al., 2015; You et al., 2015) have investigated the correlation between the expression levels of circRNAs and their corresponding mRNAs. Guo *et al.* (Guo et al., 2014; Salzman et al., 2013) and Salzman *et al.* (Guo et al., 2014; Salzman et al., 2013) found that the relative expression levels of circRNAs and their linear counterparts can differ between cell types and tissue types. Two other studies showed that the differential changes in many circRNAs are independent of the changes in the expression of their host linear transcripts upon neuronal differentiation (Rybak-Wolf et al., 2015; You et al., 2015). These results suggest that circRNAs are not simply by-products of occasional aberrant splicing (Chen, 2016). The current study provides further novel observations regarding the correlation between circRNAs and mRNAs. We found a positive correlation between circRNA expression and the expression of cognate linear mRNAs in all tissue samples (Figures 2A and 2B and Supplementary Figure S2). Furthermore, our data support the concept that circRNAs are more sensitive to changes in host gene expression. Generally, increased host gene transcription levels tend to increase the expression of both mRNAs and circRNAs; thus, if there is no splicing regulation of circRNA expression, no substantial differences in the ratio of the expression of circular and linear transcripts should be observed. However, we observed a consistently higher fraction of circular transcripts among highly expressed genes in all tissues (Figure 2C), which suggests that both transcriptional and post-transcriptional regulators contribute to the final output of circRNA expression. The correlation between the splicing efficiency of circRNAs and the total expression of their host genes may be explained by the competitive splicing between linear and circular transcripts (Ashwal-Fluss et al., 2014). Interestingly, Rybak-Wolf *et al.* (Rybak-Wolf et al., 2015)observed a negative correlation between gene expression and the logarithm of the circular to linear ratio, which is exactly opposite the trend we observed (Figure 2C). Although the cause of these paradoxical observations is currently unknown, a recent study (Liang et al., 2017a) explored how the ratio of linear and circRNA is controlled and identified many core spliceosome and transcription termination factors that control the RNA outputs of reporter and endogenous genes. It has also been suggested that circRNAs become the preferred gene output when core spliceosome or transcription termination factors are depleted. In this context, it was reasonable to observe a positive correlation, rather than a negative correlation, between the ratio of circular transcripts to linear transcripts, since the required core spliceosomes and transcription termination factors are likely to be insufficient for genes with higher expression levels. Taken together, our observations not only confirm the importance of competitive splicing against linear transcripts in circRNA production but also imply the significance of host gene transcription in determining the final output of the corresponding spliced circRNA.

Second, we investigated the relevance of both circRNAs and mRNAs for tissue-specific phenotypes. Consistent with previous studies (Ashwal-Fluss et al., 2014; Conn et al., 2015; Rybak-Wolf et al., 2015; Salzman et al., 2013; Starke et al., 2015; Szabo et al., 2015; You et al., 2015), we observed that both circRNAs (Figures 3A and 4A) and mRNAs (Supplementary Figure S3) exhibited specific expression profiles across different tissues. Our data further highlighted that the expression of these tissue-specific circRNAs was closely related to the physiological functions of the specific tissue (Figure 4B). It is widely accepted that the tissue-specific expression of linear mRNAs is related to the function of multicellular tissues and human diseases (Dezso et al., 2008; Greene et al., 2015; Melé et al., 2015; The GTEx Consortium, 2015). Therefore, it is interesting to explore the differences between circRNAs and mRNAs in terms of tissue specificity and their contribution to tissue phenotypes. In our study, we observed that the tissue specificity of circRNAs was consistently higher than that of linear mRNAs (Figures 3E and 3F). Additionally, a higher splicing ratio was observed for tissue-specific circRNAs (Figure 4C), but not for tissue-specific linear RNAs (Supplementary Figure S4), which suggests that the contribution of circRNAs to tissue specificity is somewhat independent of their cognate linear mRNAs. This finding is in accordance with the previous finding that the changes in circRNAs upon neuronal differentiation are independent of the changes of their linear counterparts (Gruner et al., 2016; Rybak-Wolf et al., 2015; You et al., 2015). Taken together with the evidence presented in previous studies (Ashwal-Fluss et al., 2014; Conn et al., 2015; Rybak-Wolf et al., 2015; Salzman et al., 2013; Starke et al., 2015; Szabo et al., 2015; You et al., 2015), we propose that while both forms of RNA transcripts can make independent contributions to tissue-specific functions, circRNAs may be more relevant to tissue specificity than linear mRNAs.

Third, we explored the accumulation of circRNAs and linear mRNAs across developmental stages in different tissues. Previous studies have shown that circRNAs gradually accumulate with age in brain samples, which has been observed in several organisms, such as humans (Szabo et al., 2015), mice (Gruner et al., 2016; You et al., 2015), and flies (Westholm et al., 2014). We confirmed a monotonic increase of circRNAs in the rat brain samples (Figure 5A), while most other tissues, such as the heart, liver, and lungs, did not present a consistent association of circRNAs with age (Figure 5A), which is consistent with the observations made by Gruner *et al.* (Gruner et al., 2016). Gruner *et al.* investigated the changes in circRNAs in mouse heart samples and found no significant changes in circRNA expression between young and aged mice (Gruner et al., 2016).

In addition to the age-dependent circRNA profile found in the brain, another major novel discovery of our study is a previously unidentified, dynamic age-dependent pattern in the testes, where circRNA levels mirror sexual maturity and the robustness of male reproduction. Our further pathway-level analysis revealed that the circRNA populations showing age-sensitive changes are essential for spermatogenesis, which was cross-analyzed and confirmed in mouse developmental stage-specific spermatogenetic cells. In particular, the potential function of circRNAs in cilium morphogenesis (essential for the formation of the spermatid flagellum) is very interesting, which may spur future extensive basic and translational studies.

Finally, we surveyed the potential causes of the age-dependent circRNA expression observed in the brain and testes. One obvious cause is the much lower degradation rate of circRN than that of mRNAs (Enuka et al., 2016), which explains the enrichment of circRNAs in exosomes (Li et al., 2015a) and several anucleate blood components, such as platelets and red blood cells (Alhasan et al., 2016). Given that neurons show relatively low proliferation, circRNAs could accumulate in neurons during aging (Gruner et al., 2016). However, increased circRNA stability cannot explain the circRNA accumulation observed in several other tissues. For example, although cardiomyocytes show similar proliferation rates to neurons, heart circRNAs do not continuously accumulate during aging (Figure 5A). Therefore, factors other than circRNA stability itself should account for circRNA accumulation during aging. Indeed, age-dependent circRNA expression has been implicated in several biological processes, including neuronal differentiation (Rybak-Wolf et al., 2015; You et al., 2015), epithelial-mesenchymal transition (Conn et al., 2015), and fetal development (Szabo et al., 2015). Therefore, it is reasonable to assume that the accumulation of circRNAs in the brain and testes is a regulatory outcome on circRNA expression to ensure proper physiological functions at different developmental stages.

In conclusion, we present a comprehensive view of circRNA expression profiles and their relevance to linear mRNAs across different rat tissues and developmental stages. We propose that circRNAs have important functional implications for tissue phenotypes and development, which are independent of their linear counterparts. Highlights of our study include the findings that the testis is the most sensitive organ showing age-dependent changes in circRNA levels and that circRNAs could be essential in regulating the process of spermatogenesis and might be employed as a biomarker of reproductive maturity and aging.

## Experimental Procedures

### Raw RNA-seq data

To explore the dynamic expression of both circular and linear transcripts based on a single dataset, we downloaded the raw RNA-seq data of the rat BodyMap dataset from the NCBI GEO database (Barrett et al., 2013) under accession code GSE53960. In the rat BodyMap dataset, Yu *et al.* (Yu et al., 2014a;Yu et al., 2014b) constructed and sequenced 320 rRNA-depleted RNA-seq libraries containing samples from 11 rat tissue types [adrenal gland, brain, heart, kidney, liver, lung, muscle, spleen, thymus, testes (male only), and uterus (female only)] from both sexes of *Fischer 344* rats across four developmental stages (2, 6, 21 and 104 weeks). For each developmental stage, four male and four female biological replicates were obtained from adrenal, brain, heart, kidney, liver, lung, muscle, spleen, and thymus tissues. For the testes/uterus, four male/female replicates were included for each developmental stage. To further investigate age-sensitive testicular circRNA expression, we also obtained raw RNA-seq data for mouse spermatogenic cells from the NCBI GEO database (Barrett et al., 2013) under accession code GSE75826. This dataset contains rRNA-depleted RNA-seq data from three different types of spermatogenic cells: spermatogonia, spermatocytes, and spermatids (Lin et al., 2016).

### Identification and quantification of circRNAs

For each sample in the rat BodyMap dataset (Yu et al., 2014a; Yu et al., 2014b), we filtered the raw RNA-seq reads by removing adaptor sequences, contamination, and low-quality reads and assessed data quality using *RNA-SeQC* (DeLuca et al., 2012). Next, we identified all circRNAs in the sample using *CIRI* (Gao et al., 2015) with default parameter settings. We constructed a rat circRNA repertoire by retaining all circRNAs with at least one back-splicing read in at least two samples of the same tissue type. After circRNA identification, we quantified the expression levels of all identified circular transcripts and known linear transcripts in the *Ensembl* rat gene annotation (Cunningham et al., 2015) (release 91) using *Sailfish-cir* (Li et al., 2017) with default settings. For each host gene, we calculated the *TPM* values of both circular and linear transcripts. The same computational pipeline was used to identify and quantify mouse circRNAs from mouse spermatogenic cells.

### Evolutionary analysis of rat circRNAs

We performed the evolutionary analysis of rat circRNAs at two levels. We compared the conservation levels of nucleotide sequences in the coding region of exonic circRNAs against those of linear mRNAs. For each exonic circRNA, we used the transcript structure of the longest linear mRNA transcript of its host gene. We downloaded the *PhastCons* scores for the rat genome from the UCSC genome browser database (Casper et al., 2018). The base-by-base *PhastCons* score represents the posterior probability that the corresponding alignment column is generated by the conserved state (Siepel et al., 2005). While a *PhastCons* score close to zero means that the site is evolutionary neutral, a *PhastCons* score close to one implies that the site is evolutionarily conserved. For each exonic circRNA, we extracted the *PhastCons* scores at all coding positions in its host gene and calculated the average *PhastCons* scores at all three codon positions for exons in circular transcripts and those exclusively in linear transcripts. In addition to nucleotide conservation levels, we analyzed the evolutionary gains and losses of rat circRNAs in the mammalian lineage. To this end, we downloaded the human circRNA repertoire from circBase (Glazar et al., 2014) and the mouse circRNA repertoire from circNet (Liu et al., 2016a). One-to-one gene ortholog tables between humans, mice and rats were downloaded from *Ensembl* (Vilella et al., 2009). CircRNAs that originated from orthologous genomic regions among different species were defined as orthologous circRNAs. We classified all the rat circRNAs with human and mouse orthologs as mammalian-common circRNAs. The rat circRNAs with mouse orthologs but without human orthologs were defined as rodent-specific circRNAs. The rat circRNAs without either human or mouse orthologs were deemed rat specific.

### Tissue specificity

To evaluate the variability of both circRNA and mRNA expression, we calculated the tissue specificity of each host gene. The *TSI* “tau” method developed by Yanai *et al.* (Yanai et al., 2005)was applied here as follows: 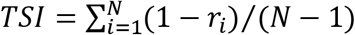, where *TSI* is the tissue specificity index; *N* is the number of tissue types; and *r_i_* is the mean expression in tissue *i* normalized to the maximum mean expression in any tissue. The *TSI* value ranges from zero to one, and a higher *TSI* implies higher tissue specificity.

### CircRNA expression-based pathway score

The *FAIME* algorithm (Yang et al., 2012) was applied to assign circRNA expression-based pathway scores for both the “cilium morphogenesis” and “spermatogenesis” GOBP terms for mouse spermatogenic cells. The *FAIME* method computes gene-set scores using the rank-weighted gene expression of individual samples, which converts each sample’s transcriptomic information to molecular mechanisms (Yang et al., 2012). A higher circRNA expression-based pathway score indicates an overall increase in the abundance of the circRNAs within the given GOBP term.

### Statistical analyses

All the statistical analyses were conducted on the R platform. PCA was used in this study to visualize the tissue specificity of circRNA expression and was performed with the “dudi.pca” function in the “ade4” library. Hierarchical clustering was conducted to visualize tissue-specific circRNAs. The corresponding heatmap was generated using the “heatmap.2” function in the “gplots” library with *Ward*’s method. The *t*-test, *Wilcoxon* test, *Spearman*’s rank correlation test, *Kolmogorov-Smirnov* test, and *Fisher*’s exact test applied in this study were performed with the “t.test”, “wilcox.test”, “cor.test”, “ks.test”, and “fisher.test” functions, respectively.

## Acknowledgements

This work was supported in part by National Natural Science Foundation of China (61372164, 61471112, and 61571109), Key Research & Development Program of Jiangsu Province (BE2016002-3), and the Fundamental Research Funds for the Central Universities (2242017K3DN04). This work was also supported by the start-up grants provided by the University of Nevada to TZ. The authors confirm that the funders had no influence over the study design, content of the article, or selection of this journal.

## Authors’ contributions

WG and TZ conceived this study. TZ, XX, ML, JS, and WG performed the analysis. TZ, XX, JJZ, KSK, TW, QC, and WG interpreted the results. WG, TZ, and QC wrote the manuscript. All authors contributed to the final version of the manuscript. All authors read and approved the final manuscript.

## Supplementary Figures

**Supplementary Figure S1. Correlation between exon numbers and the length of exonic circRNAs in rats.** The red curve represents Lowess smoothed data. The correlation coefficient (*ρ*) and *P*-value were calculated with *Spearman*’s rank correlation test.

**Supplementary Figure S2. Correlation between mean *TPM_linear_* and mean *TPM_circ_* values in rat tissues.** Each dot represents one host gene. The dots above the diagonal denote the host genes with higher circRNA expression relative to their linear transcripts. The red curve represents *Lowess* smoothed data. The correlation coefficient (*ρ*) and *P*-value were calculated with *Spearman*’s rank correlation test.

**Supplementary Figure S3. PCA of mRNA expression.** Each dot represents one tissue sample. PC1: the first principal component; PC2: the second principal component.

**Supplementary Figure S4. Fraction of linear transcripts of tissue-specific mRNAs.** The splicing ratio of tissue-specific mRNAs in the specific tissue was compared against those of all other tissue types. Each dot represents one tissue-specific mRNA. We observed a consistently lower fraction of linear transcripts for the tissue-specific mRNAs in the specific tissues (paired *Wilcoxon* test: *P* < 10^-10^).

## References

Alhasan, A.A., Izuogu, O.G., Al-Balool, H.H., Steyn, J.S., Evans, A., Colzani, M., Ghevaert, C., Mountford, J.C., Marenah, L., Elliott, D.J., et al. (2016). Circular RNA enrichment in platelets is a signature of transcriptome degradation. Blood 127.

Andergassen, D., Dotter, C.P., Wenzel, D., Sigl, V., Bammer, P.C., Muckenhuber, M., Mayer, D., Kulinski, T.M., Theussl, H.C., Penninger, J.M., et al. (2017). Mapping the mouse Allelome reveals tissue-specific regulation of allelic expression. Elife 6.

Ashburner, M., Ball, C.A., Blake, J.A., Botstein, D., Butler, H., Cherry, J.M., Davis, A.P., Dolinski, K., Dwight, S.S., Eppig, J.T., et al. (2000). Gene ontology: tool for the unification of biology. The Gene Ontology Consortium. Nat Genet 25, 25-29.

Ashwal-Fluss, R., Meyer, M., Pamudurti, N.R., Ivanov, A., Bartok, O., Hanan, M., Evantal, N., Memczak, S., Rajewsky, N., and Kadener, S. (2014). circRNA biogenesis competes with pre-mRNA splicing. Molecular cell 56, 55-66.

Barrett, S.P., Wang, P.L., and Salzman, J. (2015). Circular RNA biogenesis can proceed through an exon-containing lariat precursor. eLife 4.

Barrett, T., Wilhite, S.E., Ledoux, P., Evangelista, C., Kim, I.F., Tomashevsky, M., Marshall, K.A., Phillippy, K.H., Sherman, P.M., Holko, M., et al. (2013). NCBI GEO: archive for functional genomics data sets--update. Nucleic acids research 41, D991-995.

Bonizzato, A., Gaffo, E., te Kronnie, G., and Bortoluzzi, S. (2016). CircRNAs in hematopoiesis and hematological malignancies. Blood Cancer Journal 6.

Casper, J., Zweig, A.S., Villarreal, C., Tyner, C., Speir, M.L., Rosenbloom, K.R., Raney, B.J., Lee, C.M., Lee, B.T., Karolchik, D., et al. (2018). The UCSC Genome Browser database: 2018 update. Nucleic acids research 46.

Chen, L.-L. (2016). The biogenesis and emerging roles of circular RNAs. Nature Reviews Molecular Cell Biology 17, 205-211.

Chen, Y., Li, C., Tan, C., and Liu, X. (2016). Circular RNAs: a new frontier in the study of human diseases. J Med Genet 53, 359-365.

Conn, S.J., Pillman, K.A., Toubia, J., Conn, V.M., Salmanidis, M., Phillips, C.A., Roslan, S., Schreiber, A.W., Gregory, P.A., and Goodall, G.J. (2015). The RNA Binding Protein Quaking Regulates Formation of circRNAs. Cell 160, 1125-1134.

Conn, V.M., Hugouvieux, V., Nayak, A., Conos, S.A., Capovilla, G., Cildir, G., Jourdain, A., Tergaonkar, V., Schmid, M., Zubieta, C., et al. (2017). A circRNA from SEPALLATA3 regulates splicing of its cognate mRNA through R-loop formation. Nat Plants 3, 17053.

Cunningham, F., Amode, R.M., Barrell, D., Beal, K., Billis, K., Brent, S., Carvalho-Silva, D., Clapham, P., Coates, G., Fitzgerald, S., et al. (2015). Ensembl 2015. Nucleic acids research 43.

DeLuca, D.S., Levin, J.Z., Sivachenko, A., Fennell, T., Nazaire, M.-D.D., Williams, C., Reich, M., Winckler, W., and Getz, G. (2012). RNA-SeQC: RNA-seq metrics for quality control and process optimization. Bioinformatics (Oxford, England) 28, 1530-1532.

Dezso, Z., Nikolsky, Y., Sviridov, E., Shi, W., Serebriyskaya, T., Dosymbekov, D., Bugrim, A., Rakhmatulin, E., Brennan, R.J., Guryanov, A., et al. (2008). A comprehensive functional analysis of tissue specificity of human gene expression. BMC biology 6, 49.

Enuka, Y., Lauriola, M., Feldman, M.E., Sas-Chen, A., Ulitsky, I., and Yarden, Y. (2016). Circular RNAs are long-lived and display only minimal early alterations in response to a growth factor. Nucleic acids research 44, 1370–1383.

Fagerberg, L., Hallstrom, B.M., Oksvold, P., Kampf, C., Djureinovic, D., Odeberg, J., Habuka, M., Tahmasebpoor, S., Danielsson, A., Edlund, K., et al. (2014). Analysis of the human tissue-specific expression by genome-wide integration of transcriptomics and antibody-based proteomics. Mol Cell Proteomics 13, 397-406.

Gao, Y., Wang, J., and Zhao, F. (2015). CIRI: an efficient and unbiased algorithm for de novo circular RNA identification. Genome biology 16, 4.

Gao, Y., Wang, J., Zheng, Y., Zhang, J., Chen, S., and Zhao, F. (2016). Comprehensive identification of internal structure and alternative splicing events in circular RNAs. Nature communications 7, 12060.

Gao, Y., and Zhao, F. (2018). Computational Strategies for Exploring Circular RNAs. Trends Genet.

Glazar, P., Papavasileiou, P., and Rajewsky, N. (2014). circBase: a database for circular RNAs. RNA 20, 1666-1670.

Greene, C.S., Krishnan, A., Wong, A.K., Ricciotti, E., Zelaya, R.A., Himmelstein, D.S., Zhang, R., Hartmann, B.M., Zaslavsky, E., Sealfon, S.C., et al. (2015). Understanding multicellular function and disease with human tissue-specific networks. Nature genetics 47, 569-576.

Gruner, H., Cortes-Lopez, M., Cooper, D.A., Bauer, M., and Miura, P. (2016). CircRNA accumulation in the aging mouse brain. Sci Rep 6, 38907.

Guo, J.U., Agarwal, V., Guo, H., and Bartel, D.P. (2014). Expanded identification and characterization of mammalian circular RNAs. Genome biology 15, 409.

Hanan, M., Soreq, H., and Kadener, S. (2017). CircRNAs in the brain. RNA Biol 14, 1028-1034.

Jeck, W.R., and Sharpless, N.E. (2014). Detecting and characterizing circular RNAs. Nature biotechnology 32, 453-461.

Jeck, W.R., Sorrentino, J.A., Wang, K., Slevin, M.K., Burd, C.E., Liu, J., Marzluff, W.F., and Sharpless, N.E. (2013). Circular RNAs are abundant, conserved, and associated with ALU repeats. RNA 19, 141-157.

Kanitz, A., Gypas, F., Gruber, A., Gruber, A., Martin, G., and Zavolan, M. (2015). Comparative assessment of methods for the computational inference of transcript isoform abundance from RNA-seq data. Genome biology 16, 150.

Kumar, L., Shamsuzzama, Haque, R., Baghel, T., and Nazir, A. (2016). Circular RNAs: the Emerging Class of Non-coding RNAs and Their Potential Role in Human Neurodegenerative Diseases. Molecular Neurobiology, 1-11.

Leek, J.T., Scharpf, R.B., Bravo, H.C., Simcha, D., Langmead, B., Johnson, W.E., Geman, D., Baggerly, K., and Irizarry, R.A. (2010). Tackling the widespread and critical impact of batch effects in high-throughput data. Nat Rev Genet 11, 733-739.

Legnini, I., Di Timoteo, G., Rossi, F., Morlando, M., Briganti, F., Sthandier, O., Fatica, A., Santini, T., Andronache, A., Wade, M., et al. (2017). Circ-ZNF609 Is a Circular RNA that Can Be Translated and Functions in Myogenesis. Molecular cell.

Li, M., Xie, X., Zhou, J., Sheng, M., Yin, X., Ko, E.A., Zhou, T., and Gu, W. (2017). Quantifying circular RNA expression from RNA-seq data using model-based framework. Bioinformatics.

Li, Y., Zheng, Q., Bao, C., Li, S., Guo, W., Zhao, J., Chen, D., Gu, J., He, X., and Huang, S. (2015a). Circular RNA is enriched and stable in exosomes: a promising biomarker for cancer diagnosis. Cell Res 25, 981-984.

Li, Z., Huang, C., Bao, C., Chen, L., Lin, M., Wang, X., Zhong, G., Yu, B., Hu, W., Dai, L., et al. (2015b). Exon-intron circular RNAs regulate transcription in the nucleus. Nature Structural & Molecular Biology 22, 256-264.

Liang, D., Tatomer, D.C., Luo, Z., Wu, H., Yang, L., Chen, L.-L., Cherry, S., and Wilusz, J.E. (2017a). The Output of Protein-Coding Genes Shifts to Circular RNAs When the Pre-mRNA Processing Machinery Is Limiting. Molecular cell 68, 940-954.

Liang, D., and Wilusz, J.E. (2014). Short intronic repeat sequences facilitate circular RNA production. Genes Dev 28, 2233-2247.

Liang, G., Yang, Y., Niu, G., Tang, Z., and Li, K. (2017b). Genome-wide profiling of Sus scrofa circular RNAs across nine organs and three developmental stages. DNA Res.

Lin, X., Han, M., Cheng, L., Chen, J., Zhang, Z., Shen, T., Wang, M., Wen, B., Ni, T., and Han, C. (2016). Expression dynamics, relationships, and transcriptional regulations of diverse transcripts in mouse spermatogenic cells. RNA Biol 13, 1011-1024.

Liu, Y.-C.C., Li, J.-R.R., Sun, C.-H.H., Andrews, E., Chao, R.-F.F., Lin, F.-M.M., Weng, S.-L.L., Hsu, S.-D.D., Huang, C.-C.C., Cheng, C., et al. (2016a). CircNet: a database of circular RNAs derived from transcriptome sequencing data. Nucleic acids research 44, 15.

Liu, Y.C., Li, J.R., Sun, C.H., Andrews, E., Chao, R.F., Lin, F.M., Weng, S.L., Hsu, S.D., Huang, C.C., Cheng, C., et al. (2016b). CircNet: a database of circular RNAs derived from transcriptome sequencing data. Nucleic acids research 44, D209-215.

Melé, M., Ferreira, P.G., Reverter, F., DeLuca, D.S., Monlong, J., Sammeth, M., Young, T.R., Goldmann, J.M., Pervouchine, D.D., Sullivan, T.J., et al. (2015). The human transcriptome across tissues and individuals. Science (New York, NY) 348, 660-665.

Memczak, S., Jens, M., Elefsinioti, A., Torti, F., Krueger, J., Rybak, A., Maier, L., Mackowiak, S.D., Gregersen, L.H., Munschauer, M., et al. (2013). Circular RNAs are a large class of animal RNAs with regulatory potency. Nature 495, 333-338.

Memczak, S., Papavasileiou, P., Peters, O., and Rajewsky, N. (2015). Identification and Characterization of Circular RNAs As a New Class of Putative Biomarkers in Human Blood. PLOS ONE 10.

Meng, S., Zhou, H., Feng, Z., Xu, Z., Tang, Y., Li, P., and Wu, M. (2017). CircRNA: functions and properties of a novel potential biomarker for cancer. Molecular Cancer 16, 94.

Pamudurti, N.R., Bartok, O., Jens, M., Ashwal-Fluss, R., Stottmeister, C., Ruhe, L., Hanan, M., Wyler, E., Perez-Hernandez, D., Ramberger, E., et al. (2017). Translation of CircRNAs. Molecular cell 66, 9-21 e27.

Pollard, K.S., Hubisz, M.J., Rosenbloom, K.R., and Siepel, A. (2010). Detection of nonneutral substitution rates on mammalian phylogenies. Genome Res 20, 110-121.

Qian, Z., Liu, H., Li, M., Shi, J., Li, N., Zhang, Y., Zhang, X., Lv, J., Xie, X., Bai, Y., et al. (2017). Potential Diagnostic Power of Blood Circular RNA Expression in Active Pulmonary Tuberculosis. EBioMedicine.

Rybak-Wolf, A., Stottmeister, C., Glažar, P., Jens, M., Pino, N., Giusti, S., Hanan, M., Behm, M., Bartok, O., Ashwal-Fluss, R., et al. (2015). Circular RNAs in the Mammalian Brain Are Highly Abundant, Conserved, and Dynamically Expressed. Molecular cell 58, 1-16.

Salzman, J., Chen, R.E., Olsen, M.N., Wang, P.L., and Brown, P.O. (2013). Cell-type specific features of circular RNA expression. PLoS Genet 9, e1003777.

Siepel, A., Bejerano, G., Pedersen, J.S., Hinrichs, A.S., Hou, M., Rosenbloom, K., Clawson, H., Spieth, J., Hillier, L.W., Richards, S., et al. (2005). Evolutionarily conserved elements in vertebrate, insect, worm, and yeast genomes. Genome research 15, 1034-1050.

Starke, S., Jost, I., Rossbach, O., Schneider, T., Schreiner, S., Hung, L.-H., and Bindereif, A. (2015). Exon Circularization Requires Canonical Splice Signals. Cell reports 10, 103-111.

Szabo, L., Morey, R., Palpant, N.J., Wang, P.L., Afari, N., Jiang, C., Parast, M.M., Murry, C.E., Laurent, L.C., and Salzman, J. (2015). Statistically based splicing detection reveals neural enrichment and tissue-specific induction of circular RNA during human fetal development. Genome biology 16, 126.

The GTEx Consortium (2015). The Genotype-Tissue Expression (GTEx) pilot analysis: Multitissue gene regulation in humans. Science 348, 648-660.

Veno, M.T., Hansen, T.B., Veno, S.T., Clausen, B.H., Grebing, M., Finsen, B., Holm, I.E., and Kjems, J. (2015). Spatio-temporal regulation of circular RNA expression during porcine embryonic brain development. Genome biology 16, 245.

Vilella, A.J., Severin, J., Ureta-Vidal, A., Heng, L., Durbin, R., and Birney, E. (2009). EnsemblCompara GeneTrees: Complete, duplication-aware phylogenetic trees in vertebrates. Genome research 19, 327-335.

Wang, P.L., Bao, Y., Yee, M.C., Barrett, S.P., Hogan, G.J., Olsen, M.N., Dinneny, J.R., Brown, P.O., and Salzman, J. (2014). Circular RNA is expressed across the eukaryotic tree of life. PLoS ONE 9, e90859.

Westholm, J.O., Miura, P., Olson, S., Shenker, S., Joseph, B., Sanfilippo, P., Celniker, S.E., Graveley, B.R., and Lai, E.C. (2014). Genome-wide Analysis of Drosophila Circular RNAs Reveals Their Structural and Sequence Properties and Age-Dependent Neural Accumulation. Cell reports 9, 1-15.

Xia, S., Feng, J., Lei, L., Hu, J., Xia, L., Wang, J., Xiang, Y., Liu, L., Zhong, S., Han, L., et al. (2016). Comprehensive characterization of tissue-specific circular RNAs in the human and mouse genomes. Brief Bioinform.

Yanai, I., Benjamin, H., Shmoish, M., Chalifa-Caspi, V., Shklar, M., Ophir, R., Bar-Even, A., Horn-Saban, S., Safran, M., Domany, E., et al. (2005). Genome-wide midrange transcription profiles reveal expression level relationships in human tissue specification. Bioinformatics 21, 650-659.

Yang, X., Regan, K., Huang, Y., Zhang, Q., Li, J., Seiwert, T.Y., Cohen, E.E., Xing, H.R., and Lussier, Y.A. (2012). Single sample expression-anchored mechanisms predict survival in head and neck cancer. PLoS Comput Biol 8, e1002350.

Yang, Y., Fan, X., Mao, M., Song, X., Wu, P., Zhang, Y., Jin, Y., Yang, Y., Chen, L., Wang, Y., et al. (2017). Extensive translation of circular RNAs driven by N(6)-methyladenosine. Cell research.

You, X., Vlatkovic, I., Babic, A., Will, T., Epstein, I., Tushev, G., Akbalik, G., Wang, M., Glock, C., Quedenau, C., et al. (2015). Neural circular RNAs are derived from synaptic genes and regulated by development and plasticity. Nature Neuroscience 18, 603-610.

Yu, Y., Fuscoe, J.C., Zhao, C., Guo, C., Jia, M., Qing, T., Bannon, D.I., Lancashire, L., Bao, W., Du, T., et al. (2014a). A rat RNA-Seq transcriptomic BodyMap across 11 organs and 4 developmental stages. Nature communications 5, 3230.

Yu, Y., Zhao, C., Su, Z., Wang, C., Fuscoe, J.C., Tong, W., and Shi, L. (2014b). Comprehensive RNA-Seq transcriptomic profiling across 11 organs, 4 ages, and 2 sexes of Fischer 344 rats. Scientific Data 1, 140013.

Zhang, X.-O., Wang, H.-B., Zhang, Y., Lu, X., Chen, L.-L., and Yang, L. (2014). Complementary Sequence-Mediated Exon Circularization. Cell 159, 134-147.

